# Accurate measurement of inhibition and excitation in human motor pools during sensory stimulation

**DOI:** 10.1101/2022.09.29.509858

**Authors:** Demetris S. Soteropoulos, Alessandro Del Vecchio

## Abstract

Surface electromyography (sEMG) is a pivotal approach in clinical and basic neurophysiology, allowing us to extract the summed activity of motor units in a given muscle. Due to the bipolar nature of the motor unit action potential, the sEMG is a non-linear representation of their underlying motor unit activity and therefore affected by signal cancellation. It is not clear how this cancellation influences evoked responses in sEMG. The aim of our study was to characterise how representative an evoked sEMG response was to the firing behaviour of the underlying motor pool. To do this, we first simulated a population of motor units (and their action potentials) that responded to a stimulus with a change in firing probability. Their activity was summed then rectified to generate a simulated sEMG signal or was rectified and then summed, to generate a sEMG signal with no cancellation. By comparing the two responses to that of the underlying pool we would then compare for discrepancies. We repeated this process but by using the responses of tibialis anterior motor units to weak tibial nerve stimulation. We find that both for the simulated and experimental data the response measured through the sEMG is almost always an underestimate of the evoked response in the underlying motor pool. This is the case for both inhibitory and excitatory evoked responses. The magnitude of the inaccuracy depends on the size of the evoked response, but it cannot be accounted solely by signal cancellation, suggesting other factors may also contribute.

## Introduction

Surface electromyography (sEMG) has been a steadfast ally in the quest to understand how the brain controls movement. It allows for the non-invasive readout of the summed activity of spinal motoneurons (MNs), and offers an easy way to assess the state of the motor pool of a particular muscle, in both clinical and research settings. Despite its invaluable contributions, like any other method, sEMG has limitations and one that has been known for some time is signal cancellation. Although sEMG represents the linear sum of the signals from active motor units (MUs) in the underlying muscle (1, 2), due to the biphasic nature of MU action potentials (MUAPs), their summed activity will be an non-linear representation of their underlying activity levels (1, 3-8), as the overlap of opposing phases of MUAPs from different units will result in their cancellation in the summed signal recorded (9) from the surface of the skin. The net result is that the signal being recorded in sEMG will be an underestimate of the true level of the activity in the motor pool that innervates the recorded muscle. The degree of signal cancellation can depend on several factors, including the level of activation of the motor pool, MUAP widths, and the variability in the conduction velocity in MUs (both MNs and muscle fibres), (1, 2, 4, 5, 9). This, in turn, can influence the accuracy of measurements extracted from the sEMG (e.g.(5, 10)) although not universally so (11, 12). Inaccuracies could occur not just when measuring steady levels of contraction but also during movement or when muscles respond to a stimulus.

Evoked responses in muscles are used to assess the state of inhibitory and excitatory circuitry at various levels of the neuraxis, in both humans and animal models, using a diverse range of stimuli. Non-invasive stimulation in humans usually includes transcranial magnetic stimulation for activating cortical motor areas (13), and transcutaneous electrical stimulation of peripheral nerves for activating motoneurons and other central circuitry (see (14) for extensive references). In studies using non-human primates, electrical microstimulation is often used to quantify the connectivity of motor areas to different motor pools (15, 16). If the spiking of single neurons is available, this can be used for spike triggered averaging, where the neural spike is used as the trigger for averaging sEMG, which can help identify neurons that provide inputs to motor pools of specific muscles (17, 18). In addition, in both humans and NHPs, the state of MNs and central circuits can also be studied by measuring the responses of muscles to a mechanical perturbation (19-23). The impact of signal cancellation on the magnitude of evoked responses in sEMG though remains poorly understood and has mostly been considered when comparing the use of rectified versus unrectified sEMG for averaging (24-26).

Our first aim was to examine how much could signal cancellation potentially contribute to errors in the sEMG estimate of motor pool responses. Our second aim was to determine the degree to which this might be an issue for real data and to do this we extracted the activity of multiple MUs from the Tibialis Anterior of human participants carrying out a steady contraction during weak sensory stimulation.

## Methods

### Simulations

To examine the potential impact of MUAP cancellation on MN responses, we needed to simulate the spiking pattern of a model MN (mMN). Computationally the easiest way to do this would be to assume a Poisson Process of a desired rate, which would only require one input parameter. However, this would show an exponential interspike interval (ISI) distribution, with the highest counts at short ISIs. This would be problematic as a model for neural spiking in two ways. Firstly, due to the presence of the absolute refractory period, there are no interspike intervals <1ms in real neurons and as the membrane potential often hyperpolarises after a spike, neurons are less likely to fire another spike shortly after the previous one – this is particularly so for MNs that possess powerful afterhyperpolarisation currents (27, 28). A second limitation of using a Poisson process is that an excess of very short interspike intervals would also potentially cause cancellation of MUAPs from the same mMN, which would be particularly problematic for the purposes of this study. To be able to generate more realistic ISI characteristics, we instead simulated a simple accumulator neuron in MATLAB (Fig. 1A), whereby discreet ‘synaptic impulses’ were summated over time. The ‘membrane voltage’ (**V**) was estimated as

**Figure 1:**
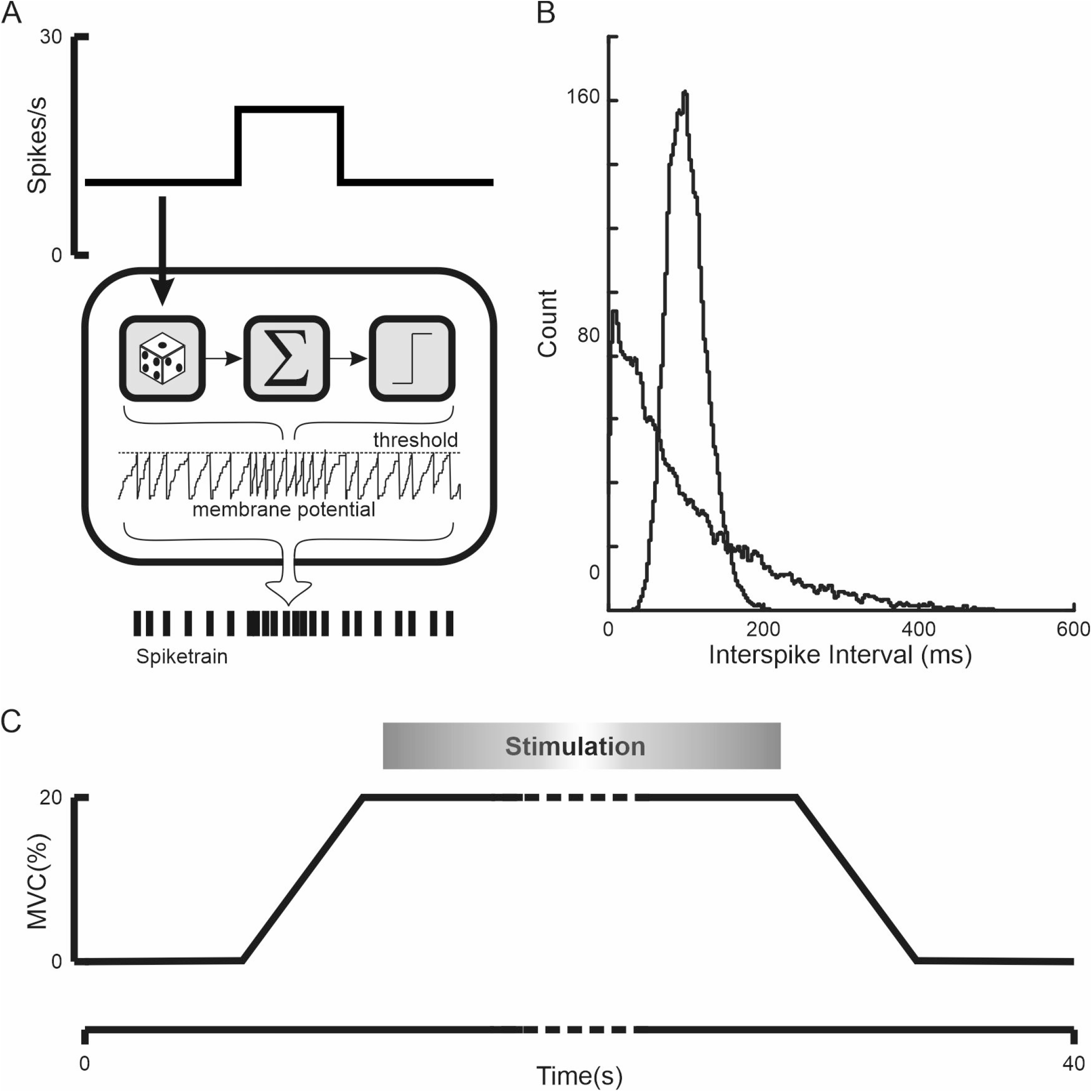
Computational and experimental methodology. A, schematic of motoneuron model used to generate spike trains of arbitrary rate modulation. The top trace shows the desired rate step profile that used to generate a sequence of inputs from a Poisson process with a similarly modulating rate. These are accumulated until a threshold is reached and a spike is assumed to have occurred. B, the interspike interval histograms of two simulated spike trains (both with a mean firing rate of 10 Hz) based on the model shown in A. By adjusting the size and rate of the inputs to the model it is possible to adjust the variance of the interspike intervals. C, Schematic of a single trial ramp for the collection of experimental data. The stimulation only occurred during the steady hold component of the trial.

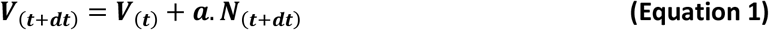

whereby **t** is time in ms, ***dt*** is a time step of 1ms, **N** is the number of inputs arriving at the mMN and *α* is the magnitude of the synaptic inputs (0< *α* <=1). The output of the mMN is a spiketrain (**S**)

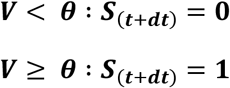

whereby *θ*is the spiking threshold (and is fixed with a value of 1). When the threshold is reached, a spike is considered to have occurred, and V resets to zero and the accumulation of events begins again – there is no membrane decay between successive time steps. The number of input events arriving at a given time step (N) are taken from a Poisson process with a mean rate of *λ* (events/ms) defined as

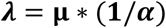

Whereby *α* is the magnitude of the synaptic inputs (Eq. 1) and **μ** is the desired firing rate of the mMN (spikes/ms). The smaller the magnitude of each synaptic input, the larger the rate of inputs to the model, and vice versa. This keeps the total input to the cell (and the output **μ**) constant for different values of *α* allowing us to control the variability of the ISI in the model independently of rate. For *α*=1, the output spike train is Poisson process, while for *α*<1, the distribution of ISIs approximates to that of a Gamma process (with a coefficient of variation of the ISIs equal to 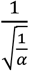) which is a more realistic fit for ISI distributions of real neurons. For the purposes of this study *α* was kept constant (*α*=0.25, corresponding to a Gamma process with regularity parameter of 4). By varying the desired spike density to the input to the mMN, we were able to generate an output rate profile of any arbitrary shape. (Fig. 1B)

### Simulated Surface EMG responses to ‘stimulation’

In order to simulate an EMG signal, we used MUAPs derived from the data collected as part of this study (described below,). This was convolved with the spike train of each mMN replacing each spike with a MUAP (Fig 2AB). These signals were then processed in two ways. In order to generate a signal analogous to a sEMG recording, the waveform mMNs of the simulated pool were summated to create a single predicted sEMG signal, which was then rectified (as would be the case for a real sEMG data, Fig. 2C). A second approach was also utilised which was to rectify first and then summate across motor units (Fig. 2DE) -this abolished the issue of amplitude cancellation but the issue of low pass filtering due to the MUAP would remain.

**Figure 2:**
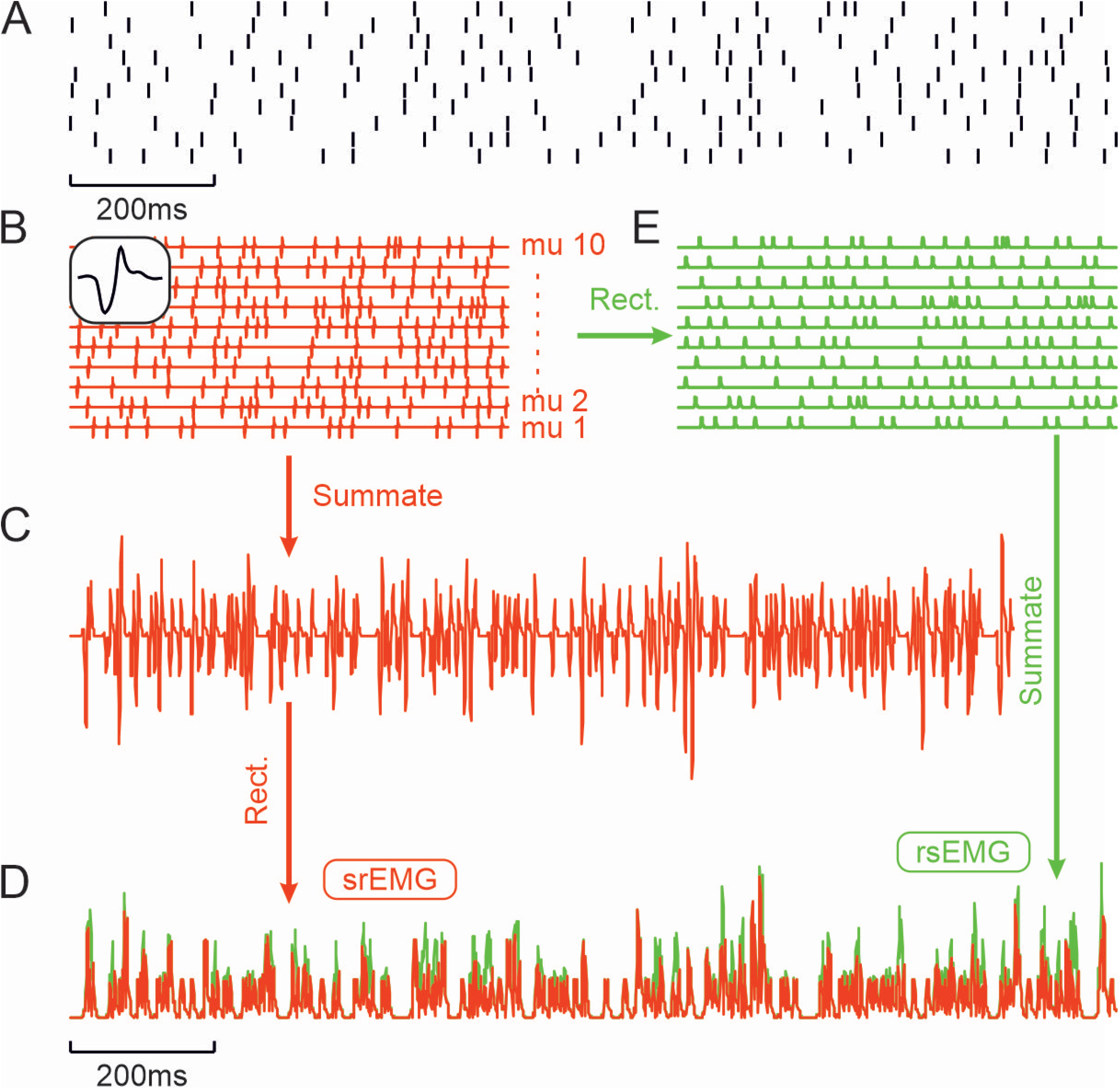
Summation of motor unit waveform data with and without signal cancellation. A, raster plot of spike train, where each row corresponds to the response of a single MU. B, This shows the result of convolving a motor unit action potential (inset) with the raster in A. C, Shows the summation of the activity of all MUs shown in C - this would correspond to the signal recorded from the surface through a bipolar EMG recording. D, in red is the rectified trace shown in C (srEMG). E, shows the result of rectifying the individual MU responses shown in B, prior to summation, which is shown as the green trace in E (rsEMG). Note the reduced magnitude of srEMG (red) compared to rsEMG(green) do to the amplitude cancellation of the action potentials prior to rectification.

The response of a motor pool to stimulation was modelled as a rate change in the responses of the mMNs (from a baseline level of 10Hz). The magnitude and duration of the responses were varied across the simulations. We assumed that 200 stimuli were given, so we repeated this process 200 times and averaged the rectified EMG and mMN responses to generate the mean EMG response and PSTH.

To assess the impact of amplitude cancellation we compared the magnitude of the simulated sEMG response to the magnitude of PSTH response. The simulated EMG and PSTH responses were first normalised by dividing by the mean value during a 100 epoch ms preceding the ‘stimulation’ period. The response was taken as the mean during from the normalised PSTH or EMG profiles across the response epoch.

### Electrophysiological Data collection

Participants provided informed consent to the study and experiments were approved by the local ethics committee at Newcastle University (Newcastle upon Tyne, UK). They were seated in a custom modified barber’s chair in a reclined position (~40° from upright), while the dominant leg was in the extended position. The ankle and foot were fastened by Velcro straps against a plate that was attached to a force transducer. Surface activity from the Tibialis Anterior (TA) muscle was recorded through a high density grid (64 contacts, 5 columns, 13 rows; gold-coated; ~1 mm diameter with 8 mm interelectrode distance; OT Bioelettronica, Torino, Italy). The skin was prepped by wiping with 70% ethanol before the grid was placed over the belly of TA and secured in place by Coflex bandage tape (3” wide) wrapped around the leg. The grid was placed with the long edge along the muscle, covering the widest point of the TA muscle. Signals were captured in a monopolar configuration relative to a reference contact placed on the kneecap. The data was continuously sampled at 10KHz / per contact, and digitised via an Intan System (RHD Recording System, Intan Technologies).

Initially, participants performed an isometric maximal voluntary contraction (MVC), and the biggest value of the rectified EMG from one of the grid contacts was used as reference for the isometric submaximal contractions (ramp contractions). The behavioural task consisted of trapezoidal ramp contractions with an increasing rate of 3.3% MVC/s (ramp duration = 6s, Fig 1C). The steady hold period lasted for 24 seconds at 20% MVC. The participants carried out 10 ramp contractions. There was a recovery time of ~1 min between ramps.

#### Stimulation

To evoke responses within the TA muscle we carried out stimulation of the posterior Tibial nerve at the ankle, during the steady hold period of each ramp. A DS7 stimulator was used to deliver monopolar stimulation (0.5ms pulse width, stimulation intensity ~2x perceptual threshold with the cathode proximal) through surface electrodes placed approximately 2 cm apart around the medial malleolus. The stimulation rate was ~ 2Hz (inter-stimulus delays of 350 to 650 ms, uniformly distributed). Stimulation was either delivered as single pulse stimuli (96 stimuli per trial) or a short train of 3 stimuli (25 trains per trial, with 3ms delay between stimuli in the train) with the two types of stimulation interspersed randomly within each trial.

#### Decomposition

During offline signal analysis, the EMG signals were band-pass filtered with a 20-to 500-Hz Butterwhorth filter and then downsampled to 2KhZ. The high-density EMG signals were then decomposed into single MUAPs by convolutive blind source separation (29). This and similar approaches have been previously validated and shows high levels of accuracy in the identification of representative populations of motor units in the tibialis anterior muscle (29, 30). The separation vector that contains an estimate of the innervation pulse trains of the motor units was visually checked and reinforced as described in (31). The motor unit waveform were then inspected visually. Only motor units showing 30 dB or SIL > 0.9 were accepted for the analysis.

## Data analysis

The extracted spike times from the decomposition were used to estimate the sEMG MUAP shape, the recruitment threshold and the response profile to the nerve stimulus for each MU.

### PSTH

The response profile of each MU to the stimulation was generated by compiling peri-stimulus triggered histogram (PSTH, 1ms bin width) of the MU spiking relative to the nerve stimulation. For the train of 3 stimuli we aligned MU activity to the time of the first stimulus in the train. The PSTH was then used to estimate the size of the responses to the stimulation for each MU. The collective response of the motor pool was estimated by adding all the PSTHs of the individual MUs.

### MUAP

The sEMG bipolar MUAP for each MU was estimated by carrying out a spike triggered average (+/-50ms relative to the spike time) of the sEMG signal (derived as described above). During the decomposition, the optimal location relative to the action potential (best signal to noise ratio) is not always the centre. For some MUs this would result in the spike time to be several ms on either side of the middle of the MUAP. To address this we shifted the spike times of each spike train to align to the centre of mass of the corresponding rectified MUAP.

## Results

### Simulation: Factors influencing accuracy of surface EMG response relative to motor pool response

Figure 3 shows results from our simulated data on how amplitude cancellation across MUs can result in an underestimation of the response magnitude in the underlying pool. In a population of 50 simulated MUs, from a baseline discharge rate of 10Hz, we imposed a 40ms change in activity ranging from 0 to 20Hz (this was the same across all MUs in the pool). The motor pool PSTH (i.e. across all MUs) is shown in Fig 3A1 showing that our model was able to generate the desired neural response profiles. Figure 3A2 depicts the mean response when the MUAPs are rectified before summating across the motor pool (rsEMG), showing that it is a smoothed (due to the duration of the MUAP) but reliable measure of the magnitude of motor pool response. However, when the MU activity is summated prior to rectification (srEMG, Fig. 3A3), which would thus potentially suffer from amplitude cancellation, the magnitude of the srEMG during the response epoch is much reduced, and this applies for both increases and decreases in rate relative to the baseline activity.

**Figure 3:**
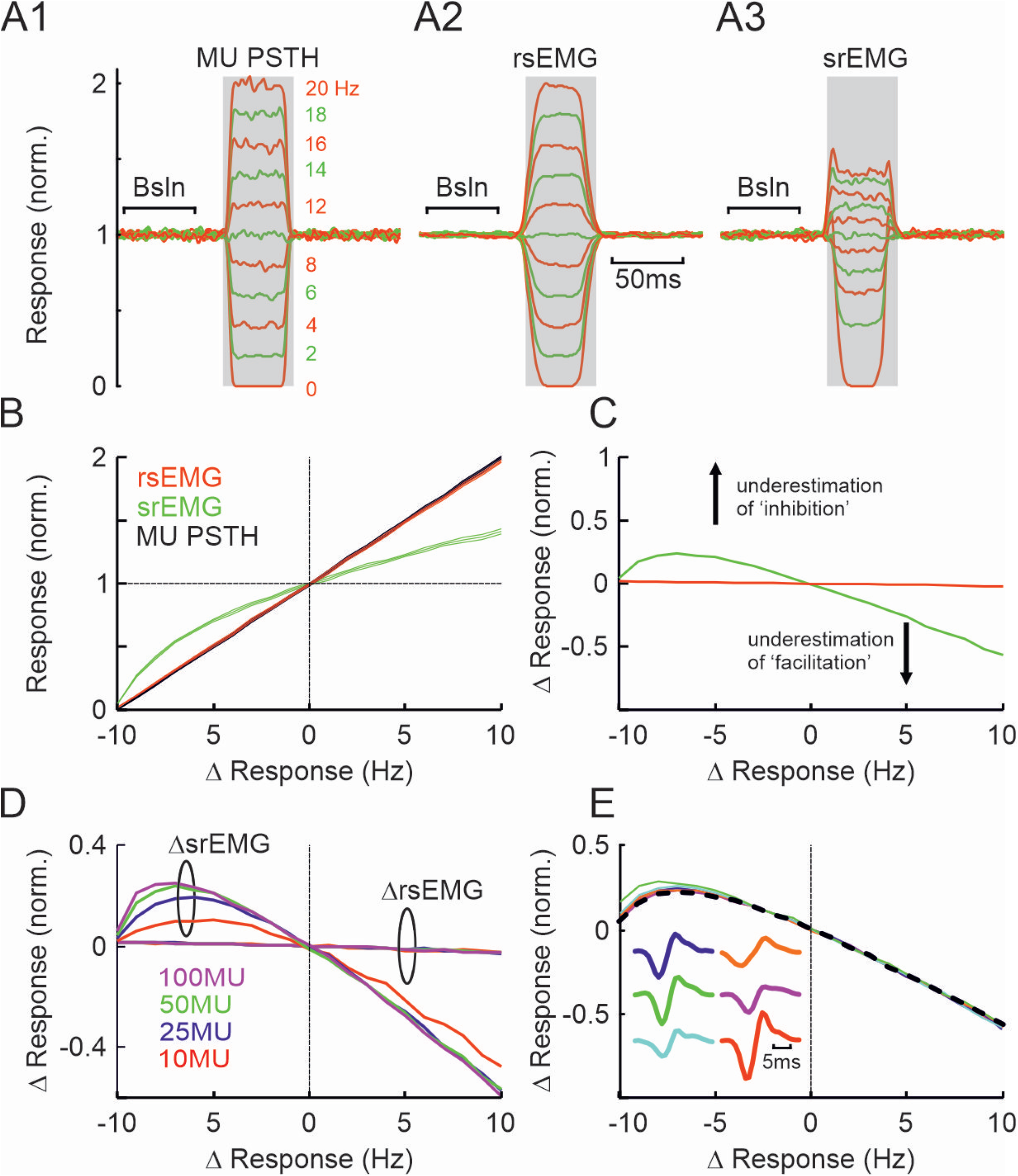
Impact of amplitude cancellation in simulated EMG responses. A, mean response of MU PSTH (A1), rectified-then-summed MUAPs (A2) and summed-then-rectified MUAPs EMG (A3). The MU population consisted of 50 MUS, with a baseline firing rate of 10Hz and a response epoch of 40ms duration. The different colours highlight the different response rate magnitudes (0 to 20Hz). The gray box shows the epoch across which estimates were made of the mean response that was used to generate the plots in subsequent panels. The responses are expressed as a fraction of mean levels during a baseline epoch (’Bsln’) of equal duration. B, this panel shows how the normalised response for the different EMGs shown in A vary versus the magnitude of the change in activity of the motor pool. The traces show the mean and 95% confidence intervals (from 20 simulations). The trace in red shows the normalised response of the simulated surface EMG with no signal cancellation (rsEMG) varies with the size of the response in the motor pool. The trace in green shows the response for the srEMG (that does suffer from signal cancellation), while the trace in black shows the normalised response of the motor pool. C, this shows the same data as in B but with the normalised response of the motor pool subtracted from the EMG responses – if there is now difference between the two then we would expect to get a flat line. D, same as C but shows the difference plot between EMG and motor pool responses when the number of contributing units to the pool was varied from 10 to 100 MUs, with each colour corresponding to a different sized pool. Note that for inhibitory responses the larger the motor pool the greater the degree of underestimation with srEMG, while this is less prominent for facilitatory responses. There is no difference in rsEMG responses. E, similar to C, but in this instance a different MUAP was used for each simulation, with the different coloured lines corresponding to different MUAP shapes. The superimposed dotted line in black is the result when each MU in the pool was assigned a different MUAP shape.

When we superimpose the response of the PSTH (black), srEMG (green) and rsEMG (red) response epochs vs the desired change in Hz of the model response (Fig. 3B), we can see that the underestimation of the motor pool response depends both on the size of the PSTH response (compare the green and black lines in Fig. 3B). For decreases in the motor pool activity relative to baseline levels, the srEMG will mirror the decrease to the same extent – this corresponds to the green line being larger than the black line in Fig 3B. This is because as the MU activity decreases, so does that degree of amplitude cancellation, but this is plotted in comparison to the baseline period of activity. For increases in firing rate, the srEMG will not increase as much -this corresponds to the green line being smaller than the black line in Fig 3B. A notable exception is when the motor pool activity stops completely in which case both srEMG and rsEMG responses reliably drop to zero. In this simulation, the rsEMG response offers a much better representation of the activity of the motor pool. Figure 3C shows a difference plot of the responses in Fig3B with the MU PSTH response subtracted from the two EMG responses. Thus for negative changes in rate (i.e a decrease in the response of the motor pool relative to a baseline period), values above zero correspond to an *underestimation* of the motor pool response. Similarly, for positive changes in the motor pool response, values below zero also correspond to an *underestimation* of the motor pool response.

Previous simulations (6) have shown that the degree of amplitude cancellation relies on the number of MUs that are active at any given time point moment. To further examine the influence of this we repeated the simulation as described above but for motor pools of different size (N=10, 25, 50, 100) and the results are plotted in Fig 3D (in the same format as Fig 3C). As the number of MUs in the pool increases, so does the srEMG underestimate of the real response, particularly for decreases in the activity of the motor pool. For increases in the response of the motor pool, the effect is similar across all motor pool sizes beyond ~10 MUs, presumably because beyond a certain number of units and rate, additional MUAPs do not cause a significant change in amplitude cancellation.

In our simulation so far we have used the same MUAP for all MUs. However real MUs will have different spike shapes and durations as picked up from the surface of the muscle, which will be determined by the actual distribution of the muscle fibres that the MN innervates and their position and distribution relative to the electrode - this could potentially influence how much amplitude cancellation takes place. To test for this we repeated the simulation as shown in Fig 3A but we used one of six different MUAPs for the simulation, with all the units in the pool having the same MUAP. The MUAPs were taken from one of the decomposed data from one of the subjects we report later in this paper. The difference plot shown in Figure 3D shows the difference plot for the srEMG responses (with each coloured line corresponding to one MUAP shape). As can be seen from the difference plot the shape of the MUAP has minimal effect on the degree of amplitude cancellation in this simulation context. As a further check we applied a different MUAP for each MU in the population and the resultant difference plot is superimposed as a black dotted line in Fig 3E, which shows the same profile.

In results there was evidence of a further non-linearity (beyond the underestimation of response magnitude) at the transition points of the rate step (see Fig 3A3 – this is most prominent for the largest deviations from baseline firing) manifesting as superimposed ‘spikes’. This is likely to be due to reduced amplitude cancellation. For the transition from low to high rate, the synchronised increase in the activity of units would mean that amplitude cancellation might be less impactful. In the transitions from high to lower firing, a similar effect is seen. In this instance the reduced amplitude cancellation would be due to decreased activity during the lower rate epoch.

To examine whether the duration of the evoked response has an impact on the sEMG amplitude we carried out a similar simulation as described above but in this instance the duration of the motor pool response was set to be much briefer (10ms, Fig 4). Figure 4A shows the mean response of the motor pool (Fig 4A1), the rsEMG (Fig. 4A2) and the srEMG (Fig. 4A3). In this instance we find that both the rsEMG and srEMG responses are quite different from the motor pool response. In the case of the rsEMG, there is now an underestimate of the response magnitude seen in the motor pool but as this cannot be due to amplitude cancellation as the signals were rectified prior to summation. This is likely to be due to the low pass filtering properties of the MUAP (see following paragraph). For the srEMG, we see, (as in Fig 3A3) non-linearities at the transition points. Of particular note is when the rate drops down to zero, there is what appears to be artifactual facilitation before and after the inhibition. The summary of the response in the EMG versus that of the motor pool at various modulation depths is shown in Fig 4B and we can now see that both the rsEMG and srEMG underestimate the magnitude of the responses in the motor pool but the underestimation in the srEMG is larger. This is shown more clearly in the difference plot of Fig4C where the srEMG line (green) deviates much more from the zero line than the rsEMG response (red line).

**Figure 4:**
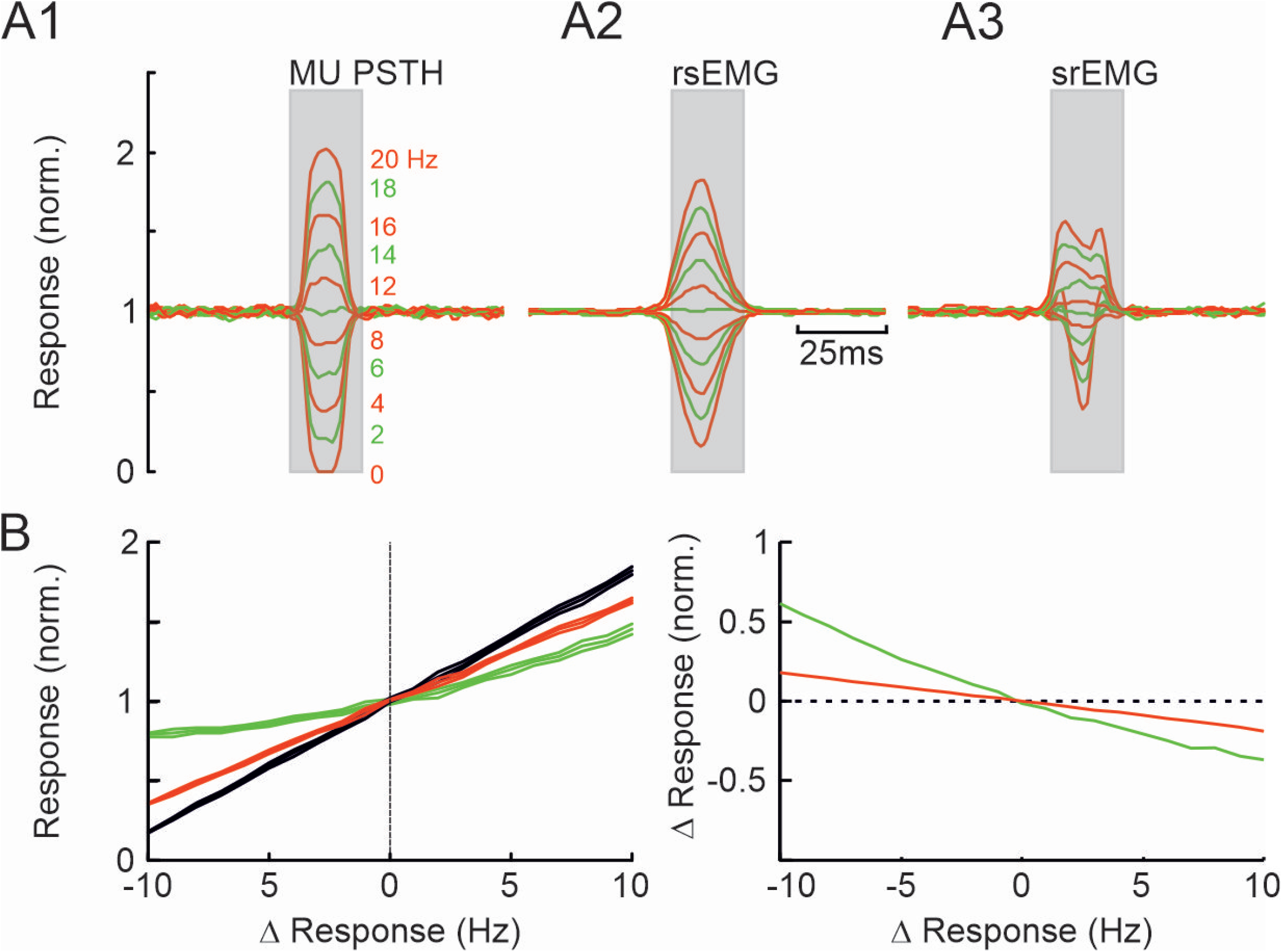
Impact of amplitude cancellation in brief simulated EMG responses. A, mean response of MU PSTH (A1), rectified-then-summed MUAPs (A2) and summed-then-rectified MUAPs EMG (A3). Simulation parameters were same as in Figure 3A except but the duration of the response on motor pool which was 10ms in duration. Note in this instance that now bow the rsEMG and srEMG responses are different to the response of the motor pool. B, this panel shows how the normalised response for the different EMGs shown in A vary versus the magnitude of the change in activity of the motor pool. The traces show the mean and 95% confidence intervals (from 20 simulations). The trace in red shows the normalised response of the simulated surface EMG with no signal cancellation (rsEMG) varies with the size of the response in the motor pool. The trace in green shows the response for the srEMG (that does suffer from signal cancellation), while the trace in black shows the normalised response of the motor pool. C, this shows the same data as in B but with the normalised response of the motor pool subtracted from the EMG responses. Note that both srEMG and rsEMG responses are now strongly deviating from a horizontal line.

### The effect of low pass filtering of the motor unit action potentials

The influence of the response duration in the motor pool is also likely to be sensitive to the duration of the MUAPs themselves. To address this, we carried a similar simulation as shown in Fig 3A but in this instance we varied the width of the MUAP used to generate the EMG response. The results are shown in Fig 5A for a response width of 40ms, and the lines of different thickness correspond to the duration of the MUAP. To generate MUAPs of different duration we resampled the real MUAP to change its width (see inset of MUAP shapes in Fig 5A, the original MUAP is indicated with a dot). As can be seen from Fig 5A2, the rsEMG response is sensitive to the duration of the MUAP and as the duration increases, so does the duration of the response while its magnitude in turn decreases – this is because the broader the MUAP the stronger the low pass filtering. This also has implications for measurements of onset latency of the response. In Fig. 5A3 we can see further evidence for non-linearities at the transition points that are also sensitive to the MUAP width – the wider the MUAP, the larger the ‘spikes’ at either end of the response regardless of whether the underlying response is a facilitation or inhibition. A similar pattern is seen in Fig 5B when the underlying response in the motor pool is much briefer (10ms). In this context the srEMG response (Fig 5B3) is dominated by the non-linearities at the response transition points. The difference plots shown in Fig 5C (for the 40ms response duration) and Fig 5D (for the 10ms response duration) show that both rsEMG and srEMG responses underestimate the response in the motor pool and this is dependent on the MUAP width – the greater the MUAP width the greater the underestimate.

**Figure 5:**
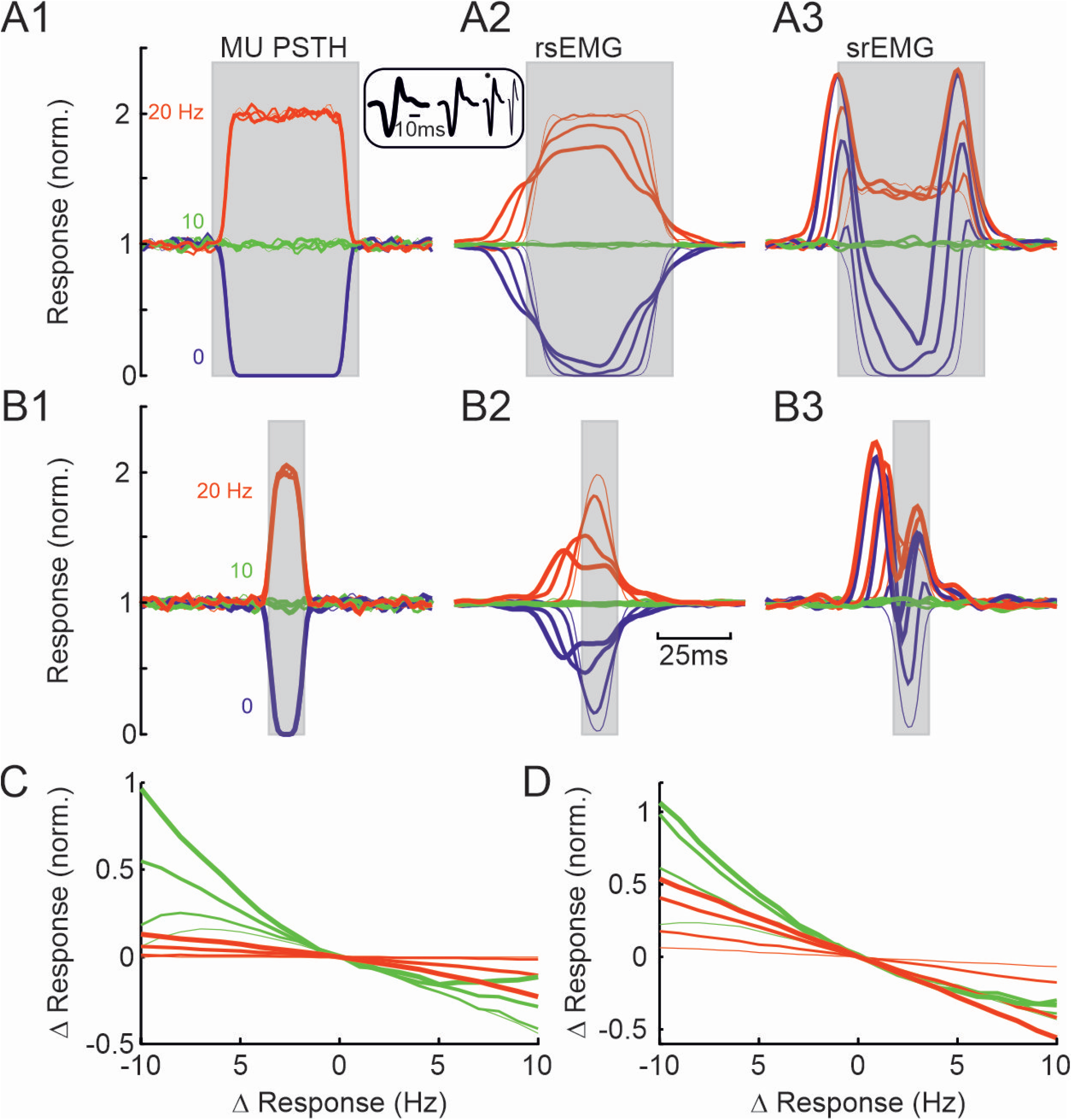
Impact of MUAP width on amplitude cancellation in simulated EMG responses. A, mean response of MU PSTH (A1), rectified-then-summed MUAPs (A2) and summed-then-rectified MUAPs EMG (A3). Simulation parameters were same as in Figure 3, but the MUAP had 4 different durations (shown in the inset, with the real MUAP indicated with a dot). The blue, green and red traces correspond to the response magnitudes of 0,10 and 20 Hz respectively, while the thickness of the traces indicate the duration of the MUAP used for the simulation. B, same as A but for response durations of 10ms (as in Fig 4). Note in both A and B that the EMG response with and without signal cancellation are different from the motor pool response. C, a difference plot between the EMG and motor pool responses show in A. Green traces correspond to the srEMG responses while red traces correspond to the rsEMG responses shown in A. The thickness of the traces indicate the duration of the MUAP used for the simulation. D, same as C but for the responses shown for the simulations with briefer responses in subplot B.

### Responses in TA to peripheral stimulation

We collected data from 6 subjects (aged 24-52, 2 female) while the carried out a ramp and hold task with the TA muscle. From those subjects we were able to extract the responses of a total of 70 MUs (range 5 to 19 per subject). Figure 6 shows the units and their responses to stimulation for an exemplar subject. Figure 6A shows the MUAP and onset of firing of the units during the ramp part of the task (during which there was no stimulation while the adjacent Fig. 6B shows the mean response of each unit to the stimulus. Figure 6C shows the population responses as recorded from sEMG (top plot) compared to the response of the decomposed motor pool (middle plot). From sEMG there a strong inhibition (minimum level of EMG is <75% relative to pre stimulus values) followed by a weaker facilitation, but in the PSTH response across all units, the suppression of activity reaches <55% of pre stimulus values. In addition, the inhibition seems to be made up of two prominent components, the earliest of which (indicated by a red arrow, beginning at ~50ms) is missing from the sEMG. Finally, the lowest plot in Fig 6C shows the fraction of units with a response larger than baseline at any given time after the stimulus – if there was no response this plot should hover around 0.5 but it can be seen that soon after 50ms most (7/10) units show suppressed firing. We can see that in this subject that, as also shown in our simulations, the sEMG response is an underestimate of what the underlying motor pool is doing.

**Figure 6:**
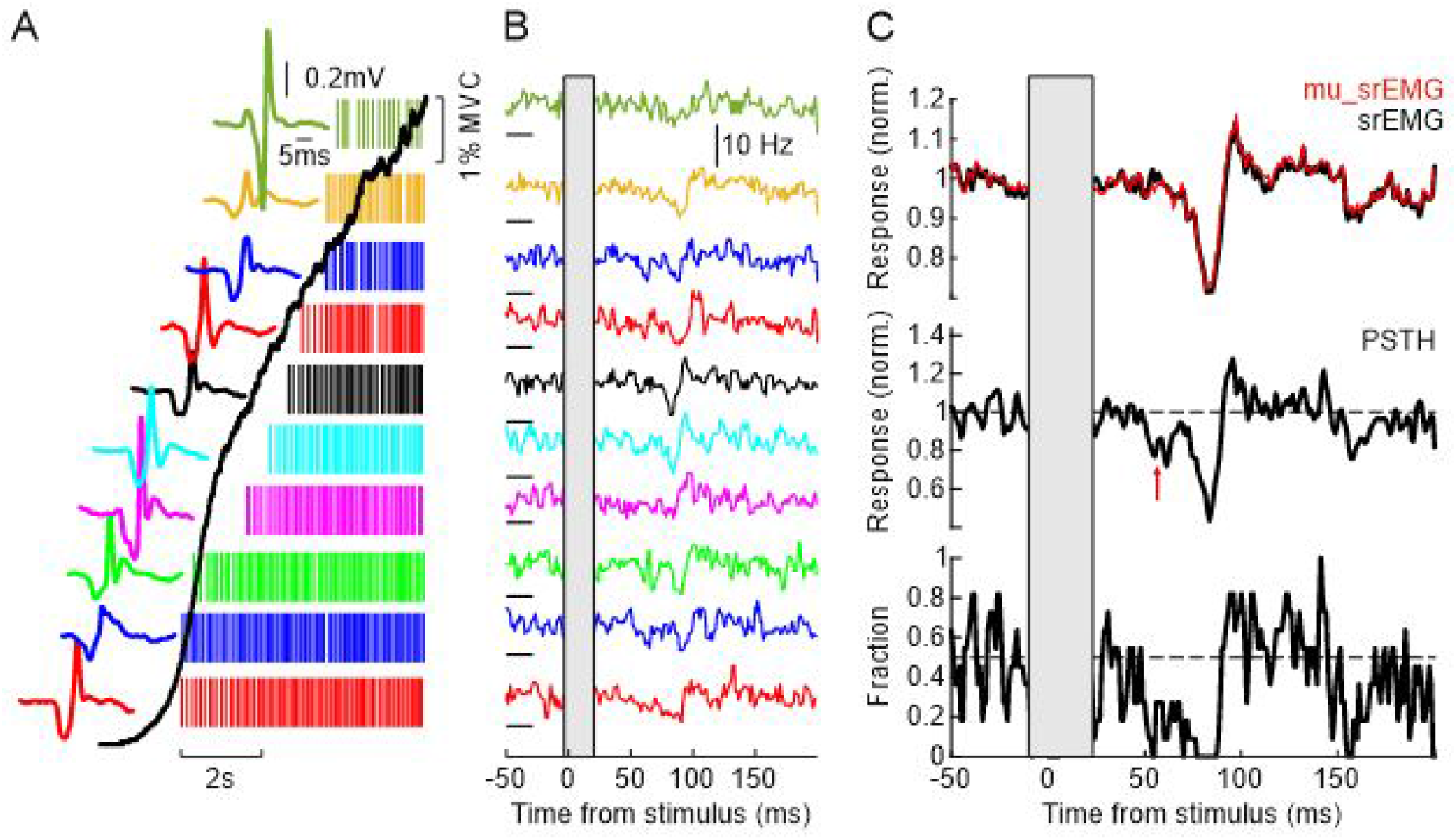
Decomposed motor units during stimulation. A, motorunit action potentials for one subjects during the ramp up of a contraction (up to 20%MVC). The raster shows the timing of the action potentials for each unit during the ramp. B, mean response to stimulation of the motorunits shown in A (colours between A and B are for matching units). The central gray bar corresponds to the epoch corresponding to the stimulus artefact. The horizontal line next to each PSTH indicates the zero level for that unit. Note the variability in the response profile for each unit. C, Top subplot is the mean response of the rectified surface EMG (sEMG) to the stimulus - note the profound inhibition between ~50-100ms. The middle subplot shows the mean response across all motor units shown in panels A and B. In both subplot the response are normalised as a fraction of levels for 50ms prior to the stimulus. The red arrow indicates a region of inhibition in the population PSTH response not seen in the sEMG response. The last subplot show the fraction of motor units that had a response larger than baseline at a given time relative to the stimulus, showing the between 50- 100ms most units showed a reliable suppression in activity. The central gray bar corresponds to the epoch corresponding to the stimulus artefact.

A caveat in this consideration is that the fraction of MU extracted through surface decomposition can vary substantially from subject to subject. As such, for real data, a smaller sEMG response might be due to a sub-total number of units was extracted and is perhaps not representative of the entire motor pool. In order to avoid this being a contributing factor we reconstituted the sEMG for each subject, but only from the decomposed MUs. The spiking of each MU was convolved with its corresponding MUAP and then summed to generate a signal similar to the srEMG used in our simulation paradigm. This is shown for an exemplar subject in Fig 7A that showed both an early excitation followed by an inhibition in response to the stimulation. The lowest trace shows the sEMG response concurrently with the PSTH response of the decomposed units. The middle trace shows the reconstituted srEMG response – this response would be susceptible to both amplitude cancellation effects but also from MUAP low pass filtering. The top trace shows the reconstituted EMG response where the MUAP for each MU was rectified first prior to summing thus removing any possible contributions from amplitude cancellation. The mean values during the inhibition showing in Fig. 7A (delineated by the vertical dotted) are shown in Fig. 7B, separately for each response. Based on sEMG the mean inhibition of EMG activity was 14% versus 26% for the PSTH. This inhibition was 20% and 23% for the srEMG and rsEMG responses respectively. Unless otherwise stated we will be primarily comparing our MU responses with the reconstituted srEMG response as this provides a more accurate (and conservative) comparison on the impact of amplitude cancellation and MUAP filtering.

**Figure 7:**
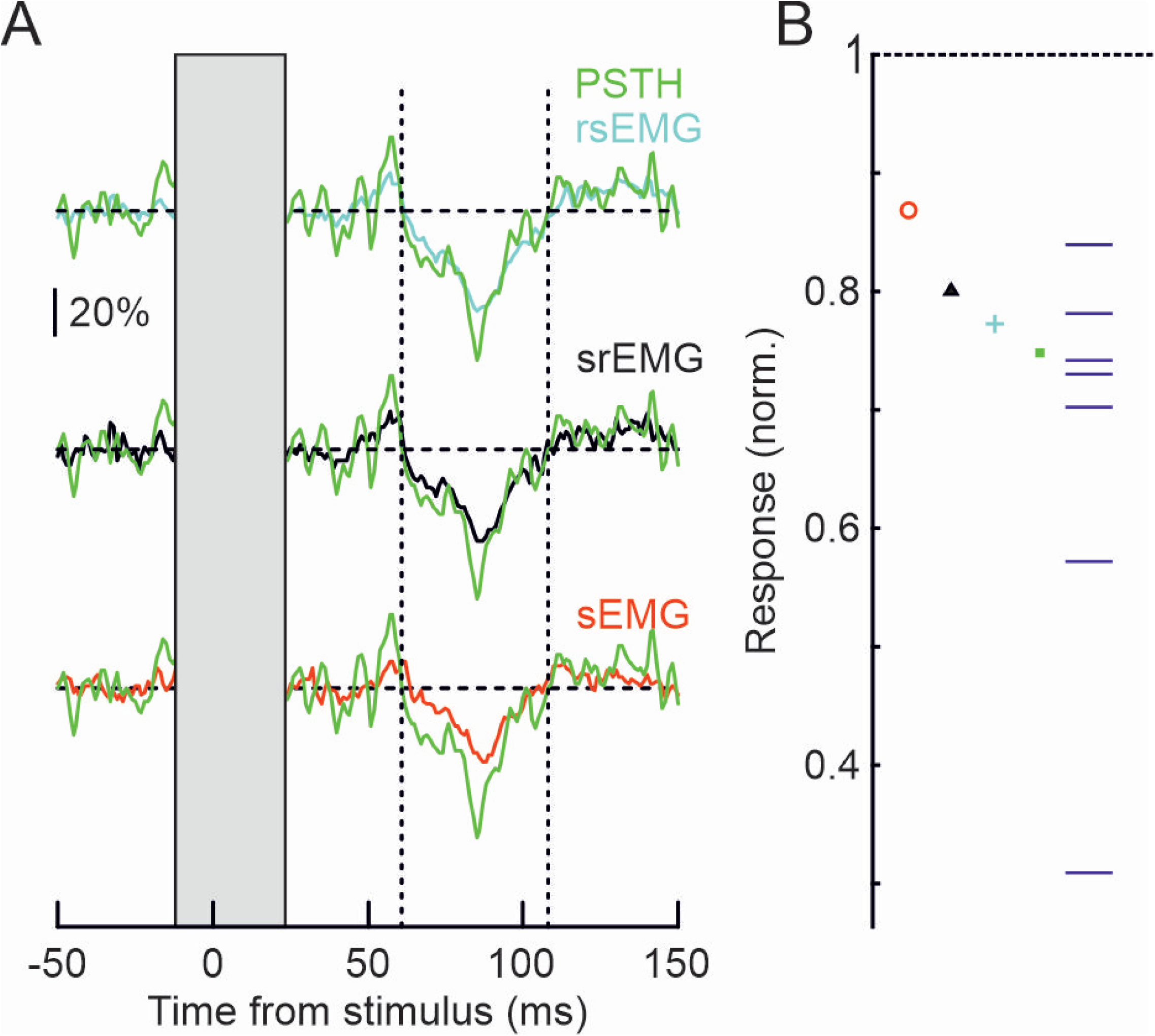
EMG responses to stimulation. A, exemplar subject rectified EMG and MU responses to stimulation. The bottom trace (red) shows the normalised EMG response recorded from the surface of the muscle (sEMG). In green is the normalised population response of the MU for this subject (and is superimposed over all EMG responses in this subplot). The middle trace (black) shows the normalised EMG response from the surface but only using the activity of the extracted MU for this subject (srEMG). The topmost trace (cyan) also shows the normalised EMG response by only using the activity of the extracted MU for this subject but in this instance the activity was rectified prior to summating (rsEMG) and hence does not suffer from signal cancellation. The vertical dotted lines delineate an inhibitory epoch during which the mean EMG levels were measured. B, this shows the mean levels of inhibition for the different EMG traces shown in A (same colour codes as in A). Note that the level of inhibition is largest in the motor pool response (green square), followed by the rsEMG (cyan cross), srEMG (black triangle) and ultimately sEMG (red triangle). The horizontal lines indicate the mean response across the motor units available for this subject.

The population results to single shock stimulation across all subjects are shown in Fig. 8, with Fig 8A showing the srEMG response for each subject. Four of the six subjects showed an early excitatory response (hereinafter referred to as ‘E1’ and highlighted with a red box) with a mean onset latency of 47.8ms (range 45 to 54ms) and mean duration of 19ms (range 13 to 29ms). All subjects showed an inhibitory response (hereinafter referred to as ‘I1’ and highlighted with a blue box). This had a mean onset latency of 69ms (range 58 to 83ms) and a mean duration of 30.8ms (range 20 to 41ms). The magnitude of the population responses (sEMG, srEMG, rsEMG and PSTH) are shown in Fig. 8B for E1 and Fig. 8C for I1 with the same colour scheme as in Fig. 7B. We also show the value of the individual MUs for each response and subject. What can be seen from Fig. 8BC is that in almost all cases, particular with the sizable responses, for both E1 and I1, the magnitude of the response in the srEMG (black triangles) was an underestimate of the underlying response in the motor pool. This can be seen collectively across the population of MUs in Fig. 8D. This shows a histogram of the difference in the size of the response (top histogram for E1 and bottom for I1) between srEMG and the individual MUs. If the srEMG was a reliable measure of the motorpool response these histograms should centre around zero, but we find that for both E1 and I1 they are both significantly different from zero. On average the E1 response in srEMG is 8% smaller than the response in motor pool (t-test, p<0.0025) while for I1 the response is 5% smaller in srEMG than in the motor pool (t-test, p<0.0001).

**Figure 8:**
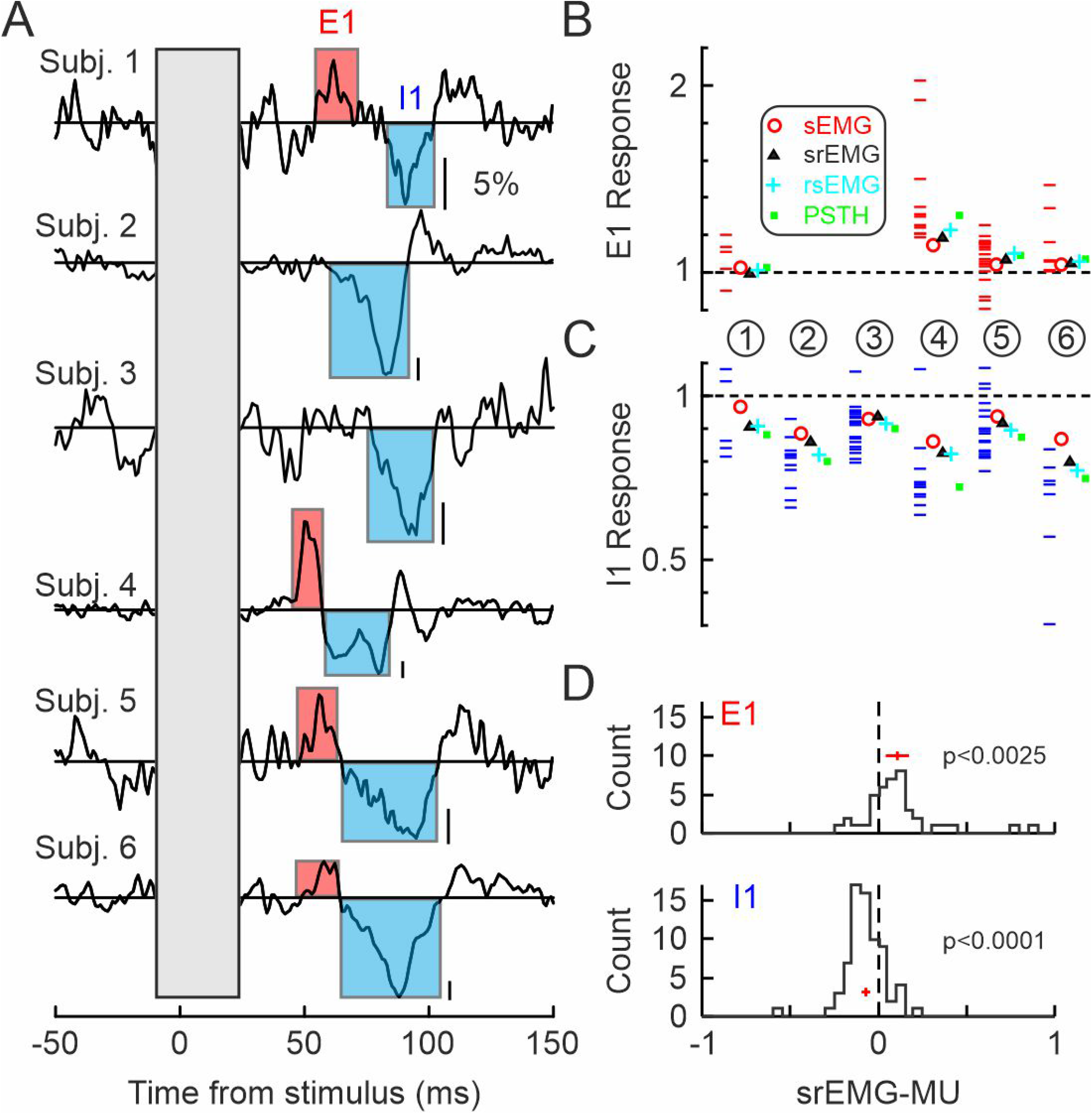
MU responses to single shock stimulation. A, srEMG responses for each subject, normalised to pre stimulus levels. Vertical calibration bars correspond to 5% of background levels of EMG. Red and blue boxes indicate early excitatory (E1) and Inhibitory (I1) epochs during which the population MU and EMG responses were analysed further. B, Mean EMG and motor pool E1 responses for each subject. Red circles correspond to the mean E1 response from sEMG, the black triangle to the mean E1 response from srEMG, the cyan cross to the response from rsEMG and the green square to the mean E1 response across the extracted motor units for that subject. The horizontal lines show the response for the individual MUs for each subject. The values corresponding to each subject is indicated by the number in the circles under each column. C, same as B but for the I1 response. D, histogram of the difference in the magnitude of E1 (top histogram) and I1 (lower histogram) of the individual MU responses with the srEMG response for the corresponding subjects. The vertical red line in each histogram indicates the mean value across all units and the horizontal red line indicates the 95% confidence intervals for the mean. For both responses there was a significant difference from zero (unpaired t-test).

We also measured the sEMG and MU responses to a train of 3 stimuli and the population results are shown in Fig 9 and follows the same format as Fig 8. Because only a smaller number of stimuli were given for this stimulation paradigm we found a smaller proportion of subjects showed a response and one subject showed no E1 or I1 response. Three subjects showed an E1 with a mean latency of 49ms (range 44 to 55ms) relative to the first stimulus in the train, and a mean duration of 17ms (range 16 to 18ms). For I1 the mean onset latency was 66.7ms (range 61 to 74ms) with a mean duration of 39.4ms (range 22 to 66ms). Figure 9BC shows the responses for each measure and MU for each subject for E1 and I1 respectively. Figure 9D shows the histograms of the differences between the srEMG response size and the MU sizes for each subject – these were significantly different for both E1, with an average 30% underestimation in srEMG vs motor pool response (t-test, p<0.007) and I1, with an average 5% underestimation in srEMG vs motor pool response (t-test, p<0.021). As with the responses to the single shock stimulation (Fig. 8) we found that the srEMG response consistently overestimated the magnitude of the response in the underlying motor pool (PSTH).

**Figure 9:**
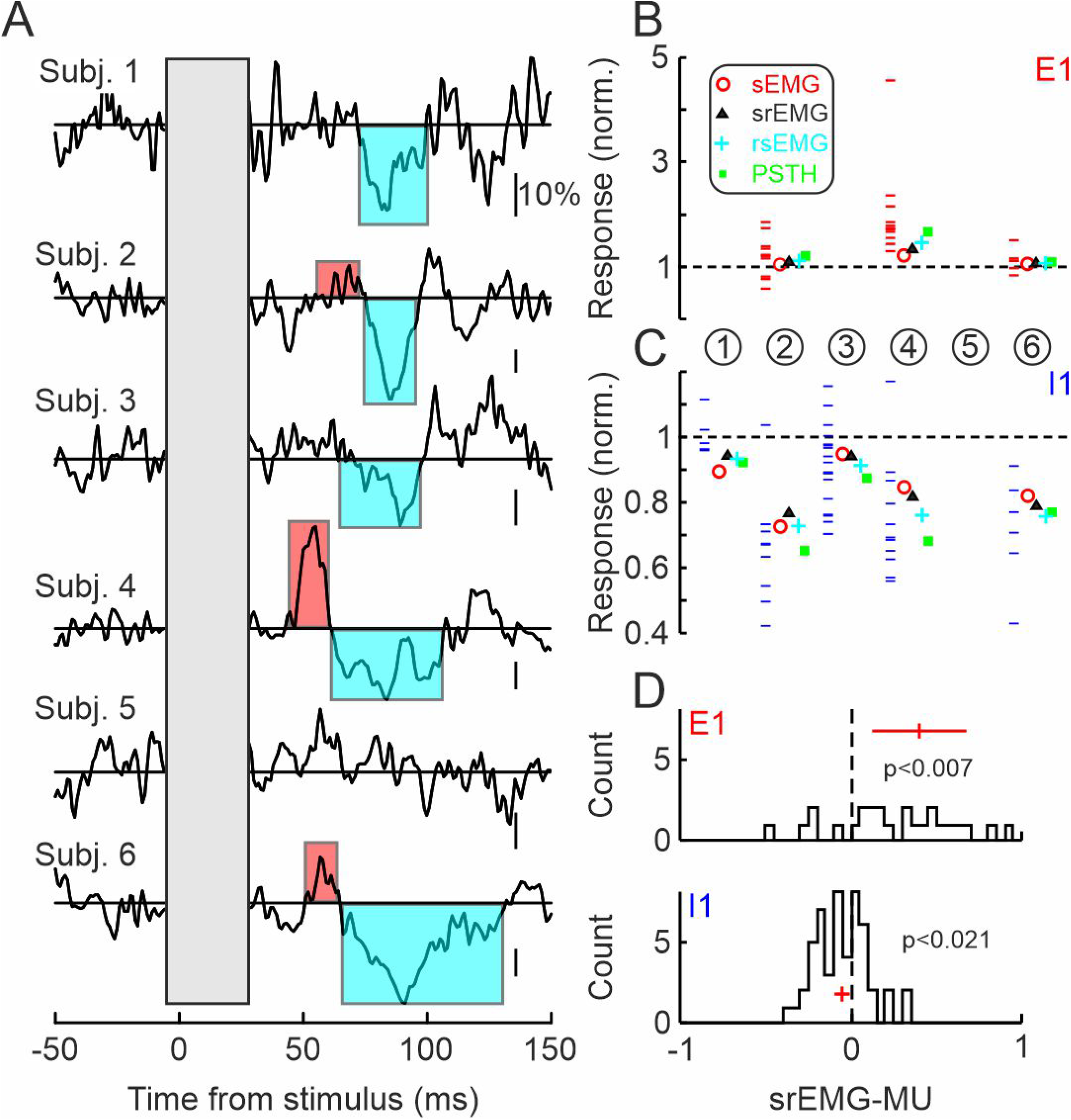
MU responses to triple shock stimulation. The format is identical to that of Figure 8 but shows the responses and population averages for triple shock stimulation

The results across both types of stimulation (single and train) and across all subjects with a significant response are shown collectively in Fig. 10. The abscissa is the net change in firing rate of the motor pool while the ordinate axis shows the difference of the normalised EMG response and the motor pool response (srEMG -MU PSTH: green, rsEMG – MU PSTH: red). The markers correspond to the responses of individual subjects while the straight lines correspond to the best fit lines for each measurement (with the same colour code as above). The linear fit was very good (R^2^>95% for both measures). The gradients of the best fit lines were significantly different from each other (non-overlapping 95% CI) and from zero. As with the simulated data we find that srEMG shows a substantial amount of underestimation of the response magnitude in the underlying motor pool and this underestimation is less when rsEMG (with no signal cancellation) is used. However even with no signal cancellation (rsEMG), there is still a significant amount of underestimation and this has to arise due factors other than signal cancellation.

**Figure 10:**
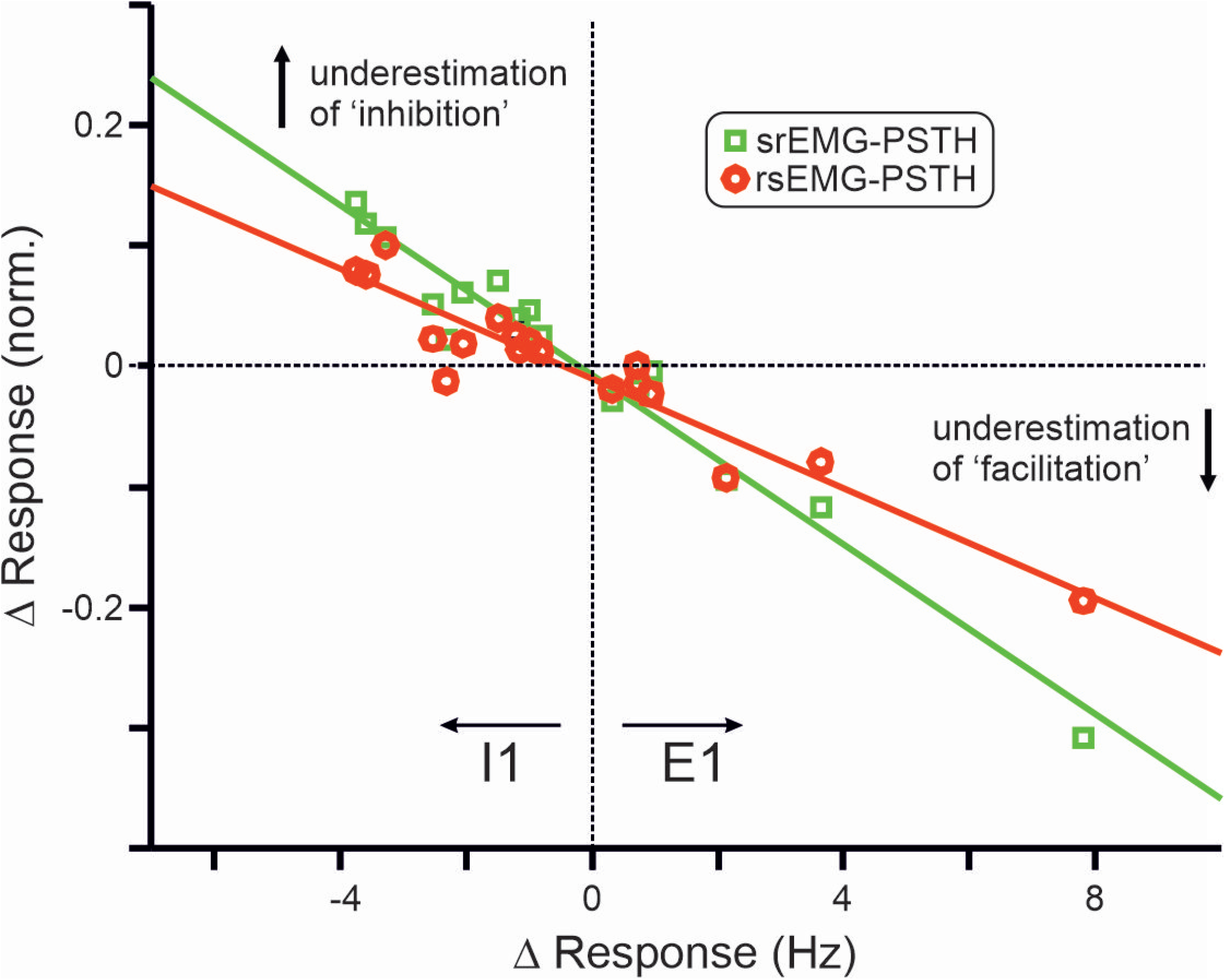
Comparison of EMG response to MU response magnitude. This shows a difference plot between the normalised magnitude of the srEMG response (green square) and rsEMG response (red circle) after the response of the MU pool has been subtracted (ordinate axis) versus the magnitude of the response in the motor pool (abscissa). Abscissa values <0 are for the I1 responses and >0 are for E1 responses. Each marker corresponds to data from one subject and responses from both single shock and triple shock stimulation are included. The solid lines show the best fit line for each response type. Note that as with simulated data the srEMG responses show a greater underestimation of the motor pool responses compared to the rsEMG, but even when using rsEMG that contains no signal cancellation, there is still a substantial amount of underestimation present.

## Discussion

This study examined how the size of the evoked response measured from surface EMG compared to that in the underlying motor unit pool. We find that, both in simulated and real data, the magnitude of the response in sEMG is an underestimate of the motor pool response. We also find that signal cancellation makes a significant contribution to this difference, although other factors are also likely to contribute. Our results indicate that this problem could likely causes inaccurate interpretations of inhibitory and facilitatory responses to different stimulation paradigms.

### Signal Cancellation

It has been known for some time that the sEMG recordings of underlying muscle activity suffer from signal cancellation – the summed waveform activity of the active MUs will be an underestimate of their true activity levels (1, 3). This means that, in turn, many measurements that rely on sEMG for estimates of MN activity and neural drive are also likely to be affected. This includes measurements of oscillatory power and coupling between muscles themselves and with other brain areas (9, 32). Moreover, the EMG amplitude cancellation increases with additional recruitment of MUs and therefore is significantly higher at high contraction forces (4-6). Therefore, the relatively synchronous MU responses after sensorimotor stimulations, may generate significantly high levels of signal cancellation and therefore inaccurate measures of facilitation and inhibition of the motor pools, as discussed below.

In addition to causing an underestimate of the magnitude of evoked sEMG response, signal cancellation can also cause the appearance of spurious effects during transitions in the activity of the motor pool, particularly when the activity of the motor pool is inhibited (33-36). In this instance a ‘pure’ inhibition can appear as a more ‘tri-phasic’ response in sEMG, being preceded and succeeded facilitatory components. At the start of the inhibition, the last components of the MUAP of the last MUs will suffer from less signal cancellation could thus contribute to peak in the rectified sEMG. Once the inhibition is over and MU activity resumes, there may also be less cancellation of the first few MUAPs (especially if they return to activity synchronously) which will also result in an erroneous peak in the rectified sEMG. We also saw evidence of this in some of our simulated data (Fig 4,5). Similarly in our experimental data we also saw evidence for features in the motor pool response not making it through to the sEMG response (Fig. 6C). The resultant sEMG response will be the net of (at least) two contrasting effects of signal cancellation – the underestimation of the magnitude of motor pool response but also the potential introduction of ‘edge’ effects, but it is unclear how common such misrepresentations are (although see (37)).

The inaccuracies we report here are unlikely to affect all stimulation paradigms equally. Those that activate MNs highly synchronously are likely to suffer much less from signal cancellation. This would include some TMS stimulation paradigms as well as M-wave and H-reflex measurements through nerve stimulation. The impact of any signal cancellation in that context would still require experimental validation though, particularly to account for the variance in the conduction velocity of MNs (38, 39) and within muscle units themselves (40). In the case of TMS, MNs can be activated indirectly via cortical and subcortical networks (41-43) which could add further potential for jitter.

Although signal cancellation contributed substantially to the underestimate the evoked responses to stimulation in both real and simulated data, alone it could not explain all of the differences between the response of the motor pool and sEMG (see Figures 4, 5, 10). Rectifying the MU activity prior to summating, which should abolish all signal cancellation, produced a more accurate measure of the motor pool response in most contexts, but this was still an underestimate suggesting other factors could contribute.

One such factor is the width of the MUAP (Fig 5) – this can vary depending on the size and distribution of muscle fibres constituting each MU but is typically in the order of tens of ms (ranging from 25 to 5ms (44)) and this means that it will also act as a low pass filter for the activity of each MU (7). Even if it was possible to reconstruct the sEMG signal from the individually rectified activity of ***<u>all</u>*** MUs in a given muscle, the response in the sEMG would be a smeared out representation of the response in the underlying motor pool due to the low pass filter properties of the MUAP, in addition to the filtering properties of the skin and other tissues between the muscle and the recording electrodes (7)-this could also have implications for measurement of onset latency of responses in sEMG versus the motor pool (see Fig 5), (but see (12)). The size and duration of MUAPs has been shown to be increased in several different neuromuscular conditions (such as motoneurone disease and spino-muscular atrophy (45)), but whether this is an issue in the measurement responses in such groups is unclear, and would require experimental validation.

### Methodological Considerations

With our simulation we sought to examine some of the factors that might influence the accuracy of the response profile in sEMG compared to that of the underlying motor pool. It was beyond the scope of this study to try to examine all possible physiological factors and their interactions, as these are likely to vary across muscles and types of contraction. It has already been shown that factors such as the synchrony between MU activity, range of MUAP size, baseline activity can affect the degree of cancellation during steady contractions (3) and the same factors are also likely to be significant during stimulation. During stimulation paradigms there is also the possibility for the contribution of MUs that are silent prior to the stimulus, thus making no contribution to the baseline activity, but that could still respond to the stimulus if it evokes a large enough synaptic input to cause the MN to fire. Similarly if the response to the stimulus differs between large and small MUs (46-48) this could also potentially influence the degree of error between surface and motor pool measures.

By extracting the activity of multiple MUs, the impact of some of these factors can be ignored or at least accounted for without direct measurement. For our experimental dataset, by knowing the activity of some of the MUs in the muscle of interest, it was possible to reconstruct what the surface EMG would look like, if these were the only units active, and measure discrepancies without having to directly measure their coupling or account for differences in baseline firing across units. Of course, decomposition of muscle activity has its own limitations and does not extract the activity of all active units during a given contraction, with an extraction bias favouring larger and superficial units(30), but at least does allow us to get a more accurate estimate of what a fraction of the active motor pool is doing.

However, even when the activity of individual MUs is known, further care is needed in interpreting their stimulus evoked responses. Particularly for polyphasic responses, secondary and tertiary modulations in activity may not necessarily arise from stimulus evoked synaptic inputs but simply occur due to the intrinsic properties of the MNs themselves – for example inhibition following an earlier excitation might simply be due to the long refractory period of MNs after a spike has occurred (49-51). Further analysis of MU activity using frequency based methods can help identify some of these potential pitfalls (50, 52, 53).

Surface EMG remains an invaluable source of information regarding the state of the muscles and their MNs but by knowing the activity of underlying MUs, the accuracy of measurements regarding the depth of inhibition and excitation of the motor can be greatly enhanced. Because of the large variety of motor units and innervation zones of the motor units in a muscle, the identification of a representative population of motor units can ultimately reveal the facilitatory and inhibitory processes that arise after sensorimotor stimulations paradigms. We demonstrate that the decomposition of EMG into constituent motor unit discharge activity into allows the precise identification of levels of inhibition and facilitation of motor pools.

## Acknowledgements

**We would like to thank the participants for their time. Funded by the MRC (MR/ W004798 /1)**.

